# Void volume fraction of granular scaffolds

**DOI:** 10.1101/2022.06.14.496197

**Authors:** Lindsay Riley, Grace Wei, Yijun Bao, Peter Cheng, Katrina L. Wilson, Yining Liu, Yiyang Gong, Tatiana Segura

**Affiliations:** Department of Biomedical Engineering, Duke University; Department of Biology, Duke University; Ninjabyte Computing; Department of Medicine, Neurology, Dermatology, Duke University

## Abstract

Void volume fraction (VVF) of granular biomaterials is a global measurement frequently used to characterize void space. There is currently no gold standard for measuring the VVF of granular scaffolds made in lab. To help the biomaterials field, we provide a library of different simulated scaffolds with known VVF for easy look-up. We use our simulated data to explore the relationship between microscope magnification and VVF, and we study the accuracy of approximating VVF using 2-D z-slice images, which is the most common approach for computing VVF of real scaffolds. Lastly, we test four different approaches for computing VVF using microscope images of real granular scaffolds and reveal that VVF may be an unreliable descriptor.

## Introduction

Void volume fraction (VVF) is one of the most commonly reported descriptors for granular scaffolds, which are biomaterials made from packed particles. The measurement describes the proportion of the scaffold that is occupied by void space between particles. In the biomaterials field, we care about understanding void space because granular materials made from microscopic particles are often designed for applications in wound healing, where cells infiltrate into and reside in the void space while they enact their therapeutic effect. VVF gives us information about particle-packing density and is one way to characterize the space available to cells. For studying the inside of granular scaffolds, the field uses light microscopy or micro-computed tomography to generate 2-D z-stack images of the sample. These slices are then either analyzed on a per-slice basis or fed into software that reconstructs a 3-D image for 3-D analysis. We are interested in studying VVF in more depth to understand how particle composition and methods of analysis affect what is being reported in the literature. Our goal is to 1) provide the field with a set of scaffold standards that can be used as a look-up chart for VVF, 2) expose the shortcomings of measuring VVF in practice, and 3) identify potential practical strategies that increase accuracy and reduce subjective variabilities.

### Background: Current approaches for imaging granular scaffolds and computing VVF

In the biomaterials field, confocal microscopy is a common imaging approach for visualizing and analyzing granular scaffolds, where particles are fluorescently labeled and/or void space is illuminated using fluorescently-labeled dextran. Scaffolds are imaged with a specified objective lens magnification (and corresponding numerical aperture) along the vertical axis to generate a z-stack of 2D-slice images that are typically a uniform step-size (z-gap) apart. In the context of reporting VVF, microscope images at 6.3x^1^, 10x^2-4^, 20x^5^, 25x^6^, and 40x^7^ magnification have been used. Protocols for z-gap size and the number of z-slices sampled also vary paper-to-paper: 1) 25-34 z-slices at a 12 µm z-gap (300-400 µm z-thickness)^2^, 2) ∼20 slices at a 5 µm z-gap (∼100 µm z-thickness)^6^, 3) 6 or more z-slices at a 17 µm or smaller z-gap (100 µm z-thickness)^8^, 4) 77 z-slices at a 1.30 µm z-gap (100 µm z-thickness)^9^, and 5) 300 z-slices at a 5 µm z-gap (1500 µm z-thickness)^1^. These variations in magnification, z-gap, and the number of z-slices can be attributed to user-preference that aims to balance image resolution, sample size, and imaging time.

Computing VVF from microscope z-stack images can be broadly categorized as 2-D or 3-D approximations, and approaches under each umbrella use techniques from image processing that introduce user bias. The most common 2-D approach is to approximate VVF by the average void area fraction among z-slices. Fiji (ImageJ) is an image processing software that approximates VVF in this way^6,8,10^, and users must choose from 16 Auto-Threshold methods for the binarization step that influences VVF^11^. The field also uses MATLAB to compute average void area fraction by implementing

MATLAB-native thresholding algorithms into custom client code, which further adds to the variability of computing VVF^2,3,9,12^. 3-D approximations first construct a 3-D volume prior to computing VVF. Imaris is a microscopy image analysis software that computes VVF by generating a triangulated mesh of either the particles or the void space, then computing the volume of the space enclosed by the mesh. User bias is introduced when setting the global threshold that is applied to all images in the z-stack during image processing. The 3-D aspect of Imaris has made it appealing for visualizing granular scaffolds as well as computing VVF^13^.

### How does particle composition influence VVF?

Particle packing theory tells us that VVF is scale-independent for homogeneous, rigid spheres, i.e., particle size should theoretically not influence VVF. However, this relationship is not always observed in real data because of image resolution issues, nonhomogeneous particles, and edge effects^12^. Therefore, we simulate a variety of granular scaffolds, referred to as particle domains, in order to observe how VVF changes with particle composition. *VVF = v / h*, and in these computational experiments, we define the scaffold’s void space, *v*, as the non-particle regions within the convex hull, *h*, of the particle centers, which helps to avoid edge effects. Our results for homogeneous, rigid spheres show a general trend of scale-independence, where the value of VVF depends on the particle packing configuration: square, hexagonal, or random (**Figure 1a**, left). As particle size increases, we see greater variation in VVF within each size category because container size is held constant at 600 × 600 × 600 µm for all particle domains, which results in fewer particles per domain (**Figure 1a**, right). With fewer particles, void space becomes significantly more variable, as we have defined it.

**Figure 1.**
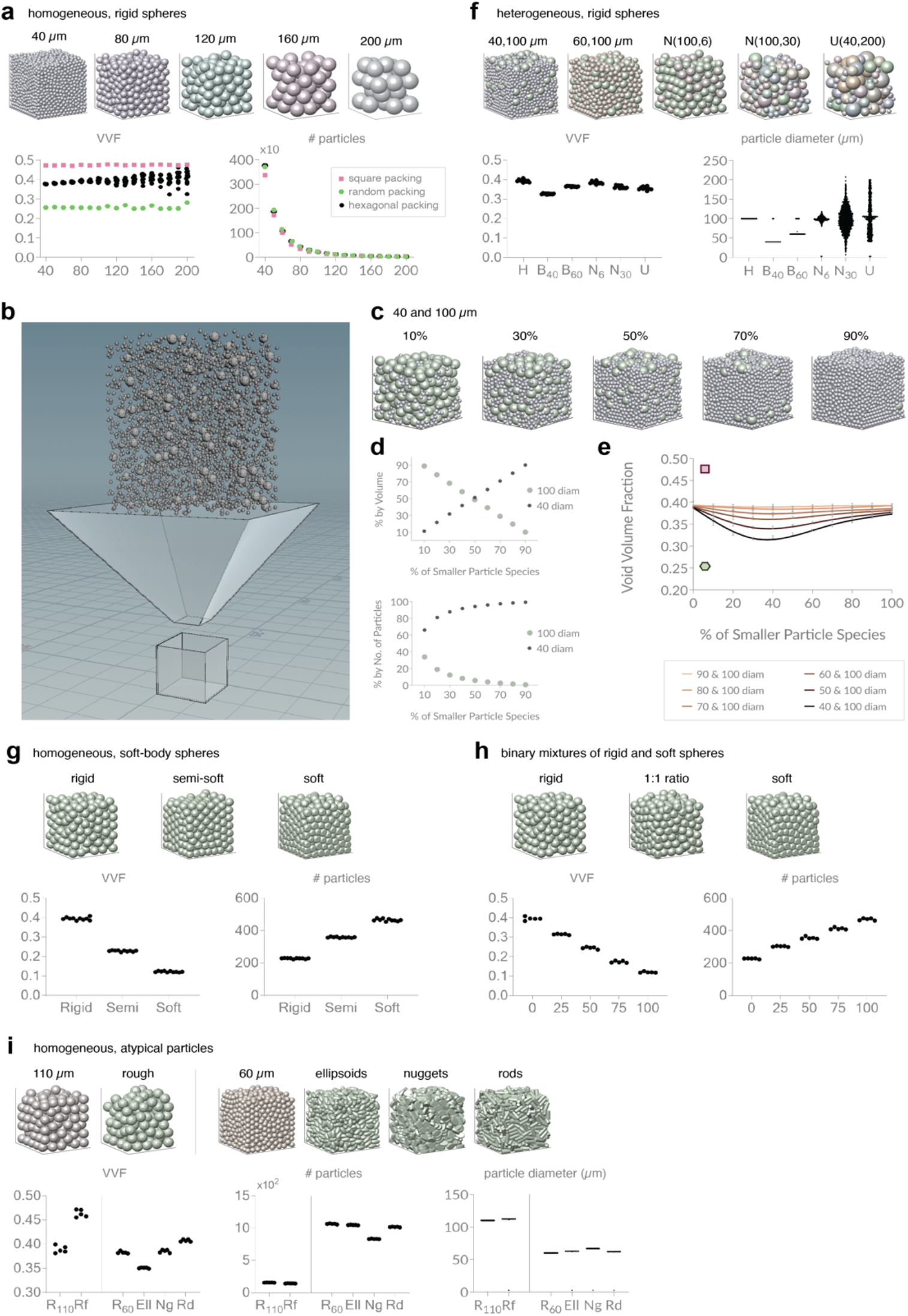
Void volume fraction (VVF) of simulated packed particles. **a**, Homogeneous, rigid spheres. Showing domain images, VVF, and number of particles in 600 × 600 × 600 µm container. X-axis is particle diameter in µm. N = 10 domains for each particle size. **b**, SideFX Houdini setup for generating binary mixtures of particles. **c**, Domain images of binary mixtures of 40 + 100 µm diameter rigid spheres by volume at varying percentages of the 40 µm species. **d**, (top) Plot confirming the percentage of 40 µm vs. 100 µm diameter particles (by volume) of our simulated domains. (bottom) Plot showing the percentage of 40 µm vs. 100 diameter particles by number of particles. **e**, VVF of different binary mixtures (see key). For each pair of particle species, we report VVF at varying percent mixtures by volume. N = 10 domains for each percent mixture. VVF of square packing (pink square) and hexagonal packing (green hexagon) shown for reference. **f**, Heterogeneous, rigid spheres. Showing domain images, VVF, and particle diameters (µm). X-axis refers to image headings; namely: H, homogeneous, 100 µm diameter particles; B_40_, binary mixture of 40 µm + 100 µm diam particles at 1:1 ratio; B_60_, binary mixture of 60 µm + 100 µm diam particles at 1:1 ratio; N_6_, normal distribution (µ: 100 µm, σ: 6 µm); N_30_, normal distribution (µ: 100 µm, σ: 30 µm); U, uniform distribution (min: 40 µm, max: 200 µm). **g**, Homogeneous, soft-body spheres. Same layout as (a). N = 10 domains for each category. **h**, Binary mixtures of rigid and soft particles. Same layout as (a). X-axis is percentage of soft particle. N = 5 domains for each category. **i**, Homogeneous, atypical particles. Showing domain images, VVF, number of particles in 600 × 600 × 600 µm container, and particle diameter (µm). X-axis refers to image headings.

We next add complexity by studying heterogeneity of particle sizes, and we find that VVF always decreases with the addition of smaller particles. We start by simulating binary mixtures with a specified proportion of large and small particles by volume. This process involves determining the number of particles according to the desired volume ratio and known container size, then initializing the particles randomly within a cylinder above the container before letting them drop into the container (**Figure 1b**). ‘Large’ particles are set at 100 µm in diameter, and ‘small’ particles range from 40 to 90 µm in diameter. We report the accuracy of our simulation methods using binary mixtures of 40 and 100 µm diameter particles (**Figure 1c,d**). Our outputs for VVF of binary mixtures of rigid spheres show that regardless of the small-particle size, VVF is minimized when around 35% of the particles (by volume) are the smaller particle (**Figure 1e**), which is consistent with the literature. Even when we increase the complexity of particle mixtures using normal and uniform distributions to sample particle diameter, we are unable to increase VVF (**Figure 1f**). Our results indicate that, for rigid spheres, VVF is maximized when particle size is homogeneous.

To study how particle stiffness impacts VVF, we generate simulated domains using soft-body physics compared to rigid body physics (see Methods). As expected, softer particles reduce VVF. This result holds for both homogeneous domains (**Figure 1g**) and binary mixtures of rigid and soft particles (**Figure 1h**), where VVF scales linearly with the proportion of soft particles. Softer particles better represent the composition of real granular biomaterials used in wound healing applications since materials are designed to mimic the non-rigidity of human tissue. As we see in our results, soft particles can produce VVF’s that dip well below the tightest packing fraction for rigid spheres, *VVF*_*t*_ = 0.26.

Lastly, we include VVF analysis of non-smooth and non-spherical homogeneous particles that reflect less common particles found in granular biomaterials. 110 µm diameter spherical particles are compared against their non-smooth (rough) counterparts, while 60 µm diameter spherical particles are compared against ellipsoid, nugget, and rod particles (**Figure 1i**). Rough particles are created by adding small bumps to the surface of smooth spheres, which results in further separation of particles from one another. VVF for rough particle domains is over 20% larger than their smooth particle counterparts, despite showing only a marginal decrease in the number of particles and a marginal increase in particle diameter (where particle diameter is defined as the diameter of the sphere with equivalent volume) (**Figure 1i**). These results show that VVF can be sensitive to small variations in particle count and size. For ellipsoid particles, our results support known findings that ellipsoids can produce tighter packing than spheres in random packing^13^. Interestingly, rod domains in random packing produce larger VVF compared to spheres and nuggets (**Figure 1i**).

### How does microscope magnification affect VVF, and how accurate are 2-D approximations of VVF?

In the biomaterials field, computing VVF of granular scaffolds typically starts with obtaining fluorescent microscope images of the sample. Imaging techniques require setting an objective lens magnification (with corresponding numerical aperture), which affects the field of view. As magnification increases, fewer and fewer particles stay in view, and therefore, a smaller sample of void space is used when computing VVF. To study how magnification influences the accuracy of VVF, we mimic scaffold magnification by cropping simulated particle domains to achieve an image-enlarging effect (**Figure 2a**). Real scaffold images at 10x, 20x, and 40x magnification are used to calibrate our crop percentages, and we maintain a similar voxel count (10^7^) for each magnification. In these computational experiments, we avoid edge effects by cropping, so we simply define void space as the non-particle regions. We simulate five different scaffolds that are cropped to 10x, 20x, and 40x magnification and report the corresponding VVF (**Figure 2b**). Our results reveal that VVF differs by up to 2% when comparing samples at 10x vs. 40x magnification. In general, the variability in VVF increases as magnification increases, which we expect considering the smaller sample size.

**Figure 2.**
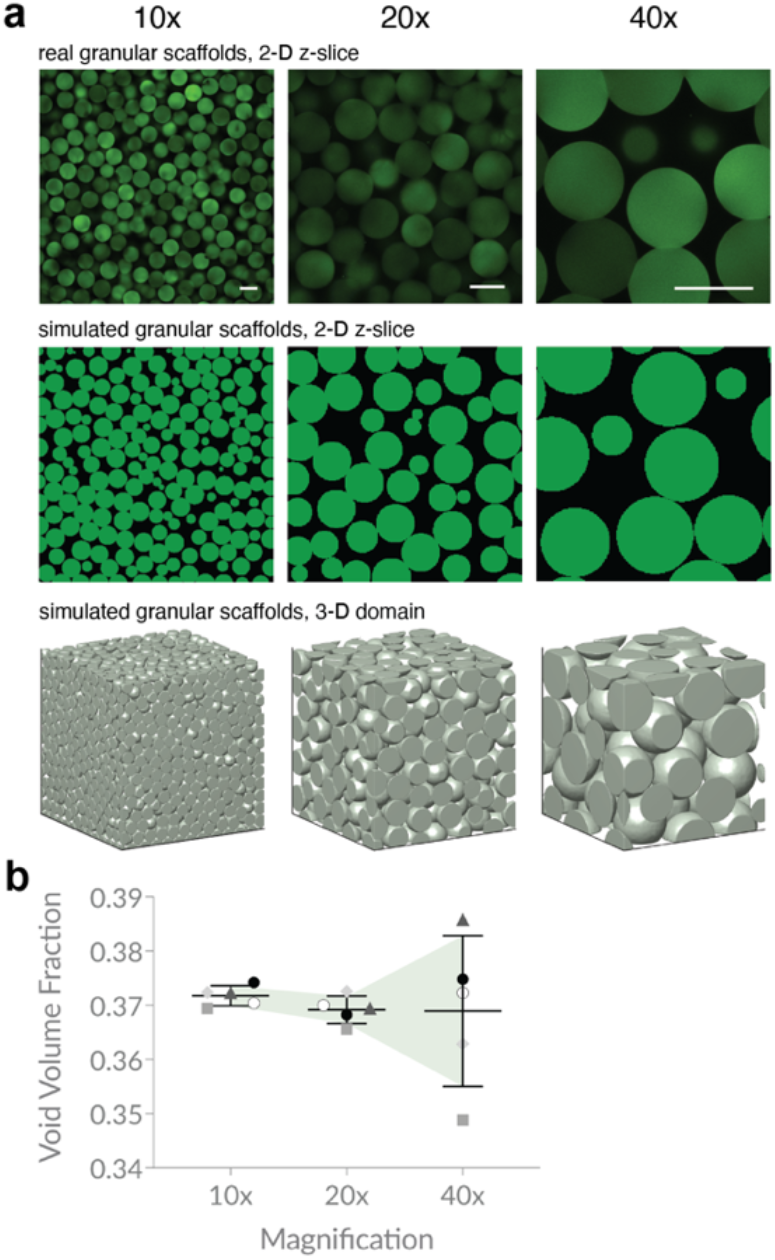
Simulating microscope magnification. **a**, Sample z-slice images from real granular scaffolds are used to calibrate our simulated scaffolds at varying magnifications. **b**, VVF of simulated domains is compared at different magnifications. Higher magnification yields less accurate VVF. N = 5 domains for each category. Scale bars = 100 µm. Errors bars denote standard deviation.

We next consider the common approach of approximating VVF of a granular scaffold by computing the average void area fraction across 2-D z-slices. We are interested in understanding how sampling 2-D z-slices influences the accuracy of the VVF approximation. To study this, we simulate five different scaffolds at three magnifications and track how VVF changes as a function of the z-gap size (**Figure 3a**). For each z-gap size, we extract the z-slices that lie within the middle half of the scaffold and that are ‘z-gap’ apart from one another, which mimics some wet lab practices aimed at avoiding edge effects. We then compute the average void area fraction among z-slices and report the accuracy of this approximation relative to the true VVF of the scaffold at the given magnification. As expected, as z-gap increases, relative accuracy decreases, and fluctuations become more dramatic. Our results show that a z-gap of less than ∼10 µm is necessary to achieve a stably accurate approximation for all magnifications, with most scaffolds converging to within 5% of the true VVF at these step sizes.

**Figure 3.**
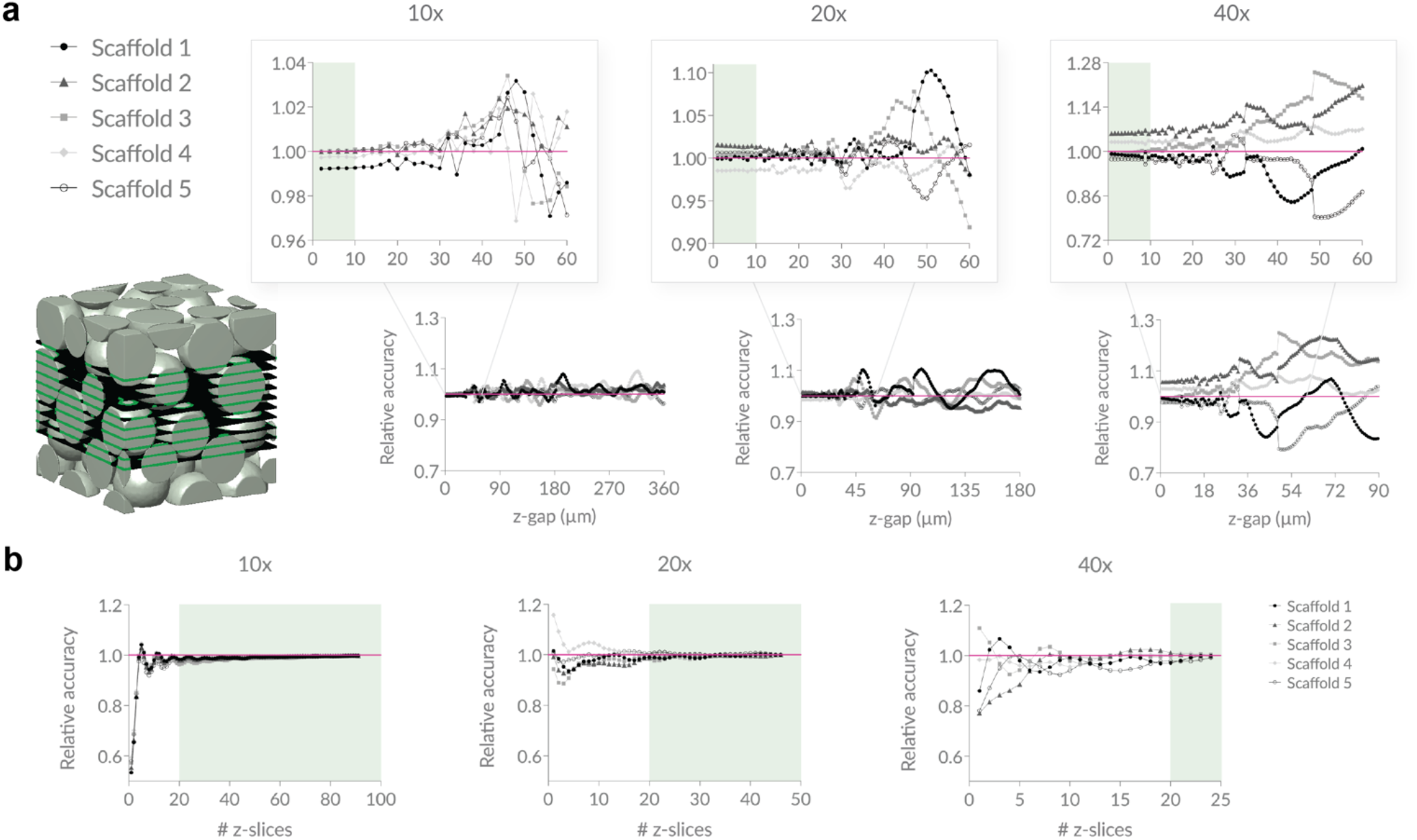
Studying z-gap and number of z-slices. **a**, Using simulated scaffolds to mimic microscope data, we approximate VVF using average void area fraction across 2-D z-slices and report how the relative accuracy of the approximation changes as a function of the step size (z-gap) taken. Z-slices are sampled within the middle 50% of each scaffold. A z-gap of less than ∼10 µm ensures a stable approximation of VVF that stays within 7% accuracy for all scaffolds across all magnifications. **b**, We again plot the relative accuracy of average void area fraction compared to the true VVF, but this time we hold the z-gap constant at 8 µm and incrementally increase the number of z-slices starting at the bottom of the scaffold. Sampling 20 or more z-slices ensures a stable approximation of VVF that stays within 4% accuracy for all scaffolds across all magnifications. N = 5 domains.

With this information, we select a reasonable z-gap of 8 µm and study how the number of z-slices influences the relative accuracy of the VVF-approximation (**Figure 3b**). Starting at the bottom of the scaffold, we sample z-slices at increments of 8 µm until the entire length of the scaffold along the z-axis has been sampled. We report the relative accuracy of the average void area fraction as a function of the number of z-slices. As expected, relative accuracy improves as the number of z-slices increases. Our results show that after ∼20 z-slices are taken, relative accuracy stays within 4% for all magnifications.

Our computational experiment indicates that both magnification, z-gap, and the number of z-slices play a role in the accuracy of approximating VVF with average void area fraction among 2D-slices. Our results are specific to our experimental setup; however, our conclusions of a z-gap less than 10 µm and at least 20 z-slices (**Figure 3**) aligns with protocols in the field used for real granular scaffolds^12^. With idealized, simulated data, we see that overall accuracy and precision of VVF improves with lower magnification because a larger percentage of the scaffold is sampled (**Figure 2b**). We next explore how magnification and the method of analysis affects the precision of VVF measurements on real microscope images.

### How sensitive is VVF to the method of analysis?

We move from simulated data to real granular scaffold data and study four different examples of software used to compute VVF. The first software is Fiji, an open-source image processing software that has built-in methods for processing microscope images and reporting void area fraction for each z-slice. The second method is simple in-house MATLAB code that uses built-in morphological operations to binarize and threshold 2-D z-slice images in order to compute void area fraction for each z-slice. The third software is Imaris, an image analysis software specifically designed for microscope images that converts 2-D slice data into 3-D surfaces renders and computes multiple measurements, including VVF. Lastly, we have developed a custom 3-D approach in Python for computing VVF that converts 2-D z-stack data into a 3-D matrix. The code segments and labels individual 3-D particles from 2-D microscope z-stacks, which can then be used to extract VVF. We refer to this method as 3-D Particle Segmentation.

All software that analyze real images implement techniques from image processing to classify void space versus non-void-space pixels, and these steps require user input. We are interested in studying the variability of VVF that can arise from a reasonable range of user-inputted threshold parameters or settings. For each software, we determine a user-inputted range that generates a ‘low,’ ‘medium,’ and ‘high’ version of the output image, where the categories represent the relative number of void space pixels in the final binarized image. Specifically, low output images show many ‘false particle’ regions but at least one ‘false void’ region; middle output images show both ‘false particle’ and ‘false void’ regions; and high output images show many ‘false void’ regions but at least one ‘false particle’ region.

To study VVF of granular scaffolds made in lab, we have chosen to focus on fluorescently-labeled particles as opposed to fluorescently-labeled dextran (which illuminates the void space) in order to take advantage of software developed by our lab. Both labeling options are conceptually the same. We study two different microporous annealed particle (MAP) scaffolds, which are a specialized type of granular scaffold comprising interlinked hydrogel microparticles (HMPs)^7^. The first MAP scaffold contains ∼70 µm HMPs made of modified polyethylene glycol^14^, and the second contains ∼100 µm HMPs made of modified hyaluronic acid^15^. For each piece of software, we analyze 10x, 20x, and 40x images of each scaffold and plot VVF based on input parameters that result in images classified as low, medium, and high (**Figure 4**). Below each software type, we show representative images from each magnification, as well as the corresponding binarized image output for our three categories. We see notable variability among different types of software, among different magnifications, and even among the reasonable range of user-inputted settings. For each scaffold, we also include a summary plot that compares the middle-range VVF values for all methods at each magnification (**Figure 5**). The trends indicate that results are scaffold-specific. Our second sample scaffold contains larger, brighter particles relative to the first scaffold, which contributes to the narrower VVF range in the low, medium, and high categories. Not only do these results demonstrate that precision is greatly affected by image quality, but in addition, regardless of image quality, our results reveal discrepancies among different magnifications, as well as different software. For example, Figure 5b shows a statistical range in VVF of 15% among software outputs for 10x magnification.

**Figure 4.**
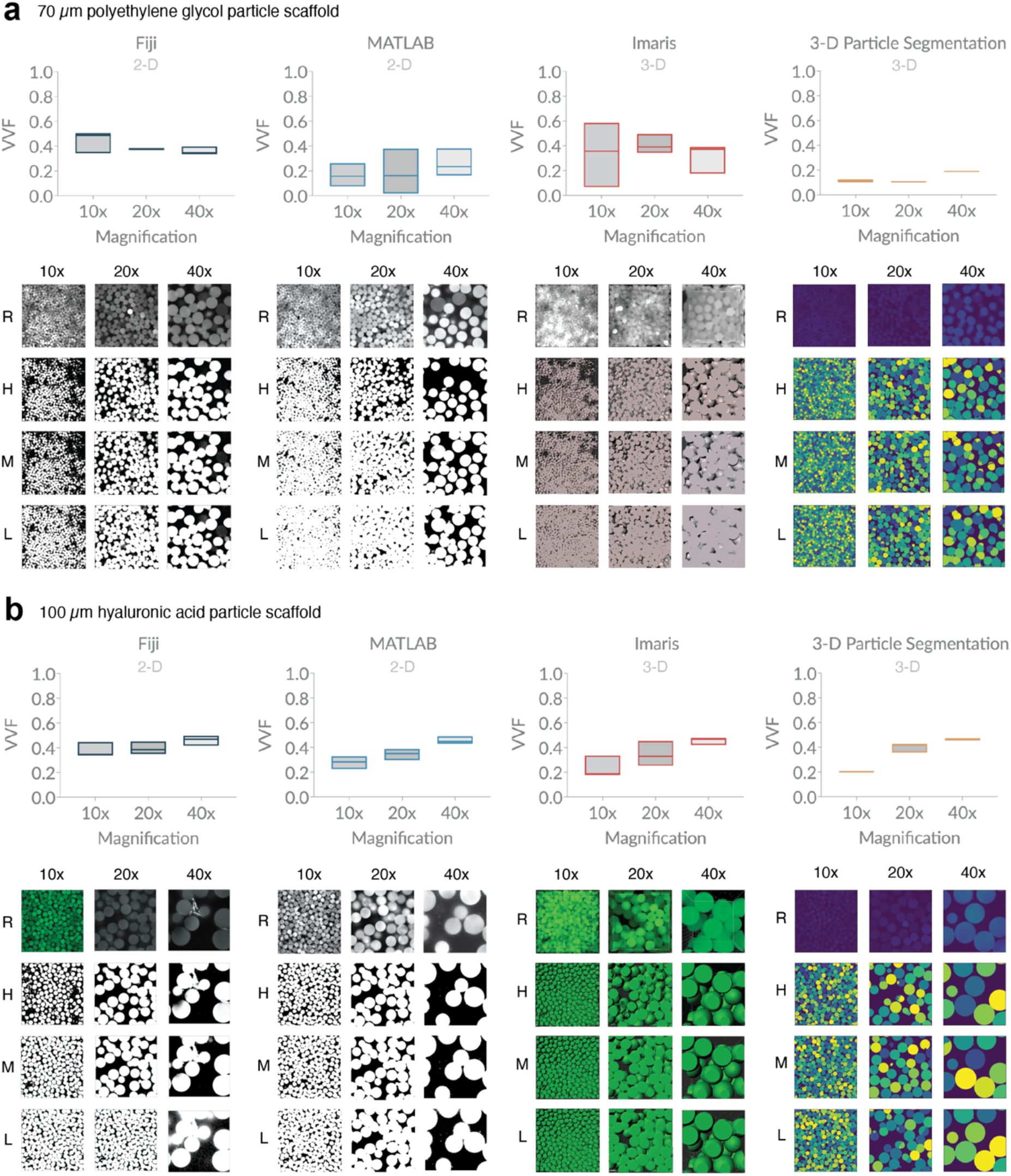
Variation in void volume fraction (VVF) of granular scaffolds across different software, magnification, and user-bias. **a**, Scaffolds comprising 70 µm diameter polyethylene glycol (PEG) particles were analyzed for VVF using Fiji, a custom MATLAB code, Imaris software, and a custom particle segmentation code. For each magnification, we report box plots indicating a low, middle, and high VVF that reflect variation in user-inputted parameters or settings. Below each software plot, we show sample z-slice images of real microscope images (R), as well as high (H), middle (M), and low (L) binarization outputs. Scaffolds were imaged with the following z-gap and number of z-slices: (10x) 4.075 µm, 36 slices, (20x) 1.20 µm, 118 slices, (40x) 1.00 µm, 142 slices. b, Scaffolds comprising 100 µm diameter hyaluronic acid (HA) particles were analyzed in the same manner as (a). Z-gap and number of z-slices were as follows: (10x) 3.875 µm, 40 slices, (20x) 1.20 µm, 101 slices, (40x) 1.10 µm, 192 slices.

**Figure 5.**
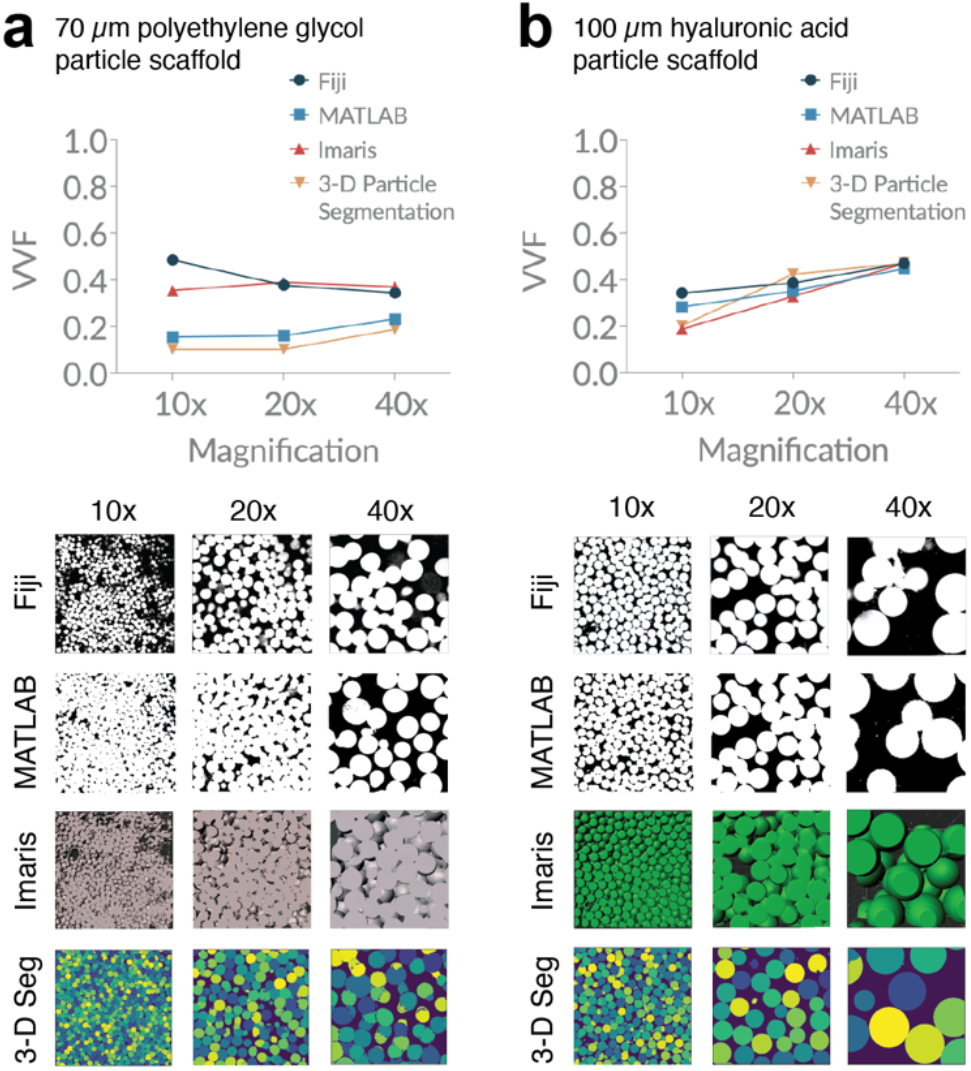
Microscope magnification and analysis software influences VVF measurements. **a**, Our PEG scaffolds highlight the variation in VVF that can be seen across different software when ‘middle’ range parameters and settings are used. **b**, Our HA scaffolds highlight the variation in VVF that can be seen across different microscope magnifications. Sample z-slice images are shown below for comparison.

We unfortunately have no way of testing the accuracy of the outputs since no gold standard exists, but our results are striking because they reveal how the most commonly-reported scaffold measurement of VVF may, in fact, be subject to substantial variance. At minimum, they reveal that VVF is highly dependent upon parameters used in their computation, including microscope magnification, the software being used, and user-inputted threshold settings. We generally recognize that when imaging granular scaffolds, 40x magnification produces sharper particle borders, which makes void space delineation more accurate. **Figure 2b** indicates that 40x magnification shows the greatest variation in VVF relative to lower magnifications; however, the relative percent difference of ∼10% suggests decent precision among samples. Therefore, sampling several regions of interest at 40x magnification across the scaffold and reporting the average VVF should give a reasonable approximation to true VVF.

## Conclusion

In the biomaterials field, void space characterization is often summarized with a single reported value of VVF. Since there is currently no gold standard for measuring VVF of real granular scaffolds, we have created a vast library of ideal granular scaffolds using simulated data, and we report the VVF of each scaffold to serve as a resource for the scientific community. In practice, VVF is computed from microscope z-slice images; however, the accuracy of VVF is highly dependent upon multiple factors, including microscope magnification, the region of interest, z-slice sampling, z-intensity corrections, the software used to compute VVF, and user-selected settings. To address these issues, we analyze both ideal and real data.

Using ideal, simulated scaffolds provides us with the true VVF by which to compare approximations. We first use simulated scaffolds to study how microscope magnification influences the accuracy of VVF, and we find that higher magnifications produce more inaccurate approximations. Next, we use simulated scaffolds to study how the microscope step size (z-gap) and the number of sampled z-slices influence the VVF measurement. Based on our results, we recommend using a z-gap of less than 10 µm and at least 20 z-slices for the optimal sampling.

To study how software choice affects VVF measurements, we use four different software options to analyze microscope images of real granular scaffolds at different magnifications. We range user-inputted parameters or settings to obtain a reasonable range of VVF outputs for each software, and our results highlight the variation in VVF that can be attributed to magnification, software, and user-bias when selecting parameters. Sampling multiple regions of interest at the highest magnification may help to obtain the most accurate VVF for real granular scaffolds.

While it is important to be mindful of these nuances when computing and reporting VVF, our findings also highlight the need for more quantitative descriptors for characterizing the void space of granular scaffolds.

## Methods

### Simulating granular scaffolds and computing VVF

Packed particle domains were simulated using SideFX Houdini software. Particles were randomly initialized above a funnel geometry that feeds into a 600 × 600 × 600 µm container. Domains were discretized on a uniform Cartesian grid, where 1 grid unit = 1 µm. The mesh size, *dx*, for all domains was chosen to produce a total voxel count between 10^7^ and 10^8^, which produced the best trade-off between detail and memory usage. For rigid particles, we use Houdini’s native rigid-body physics solver to simulate how rigid particles fall, collide, and ultimately settle in the container. Domains containing binary mixtures are generated by first determining the number of species 1 and species 2 particles according to the desired volume ratio and container size, then initializing their starting positions (randomly) within a cylinder above the setup. For non-rigid particles, we use Houdini’s native finite element physics solver to simulate non-rigid particles, which is a well-known technique for simulating soft-body physics. We generate ‘semi-soft’ and ‘soft’ spherical particles by adjusting the Lamé parameters of the material model used in the finite element solver. Semi-soft particles have Lamé’s first parameter, λ, equal to 1000 and Lamé’s second parameter, µ, equal to 250, while soft particles have λ = 400 and µ = 50. For rough particles, we use Houdini’s modeling capabilities, where points are randomly scattered on the surface of a sphere to be used as sources of bulging for creating the final effect. We then add bumps to 100 µm particles, and bumps are approximately 3 µm tall. Lastly, we use Houdini to model atypical particle shapes, such as ellipsoids and cylinders (rods). Nuggets were created by extruding a parametric curve that describes the perimeter of an egg-like shape. Ellipsoids are 100 µm along the major axis and 50 µm along the remaining axes. Nuggets are 100 µm long, 65 µm wide, and 30 µm thick. Rods are 100 µm long and 40 µm in diameter. Isotropic ellipsoid domains were generated by first initializing isotropic ellipsoids in pseudo-hexagonal packing inside the simulation container, then allowing them to settle using Houdini’s rigid-body solver.

For homogeneous, rigid, spherical particle domains, we only include spheres that fit entirely within the boundaries of the container. For all other domains, we crop particles at the height of the container, i.e., 600 µm in the z-direction. Once a domain has been simulated, the particles are rasterized to a uniform 3-D grid. Since the domains that we generate range from rigid spheres to non-rigid custom geometry, a general data format is required. We use a map data structure to store information about which grid voxels belong to each particle, where each particle is uniquely labeled with an integer ID. This data structure is then written out to a JSON file, along with any other fields one might need for analysis, such as, but not limited to: domain size, total particle count, total voxel count, voxel count per particle, and voxel size (mesh size). For rigid spheres, a simple CSV file storing the particle centers and radii, as well as the mesh size, is used.

### Simulating magnification and studying z-slices

We use microscope images of real granular scaffolds comprising 100 µm diameter particles as our reference z-slice images for 10x, 20x, and 40x magnification. To start, we calibrate our simulated domains to 10x magnification by using an 800 × 800 × 800 µm domain of 60 µm spherical, rigid particles and cropping inward 4% in all directions, which produces 736 × 736 µm 2-D slice images. We opt for 60 µm particles instead of 100 µm particles to avoid simulating particle domains that exceed 800 × 800 × 800 µm. This initial crop is to avoid edge effects; however, the effect of particle alignment against the walls of the container is still present, e.g., Figure 3b, for small z-slice numbers, there is a dip in relative accuracy for all scaffolds at 10x magnification; Figure 2b, the average VVF at 10x is slightly larger than 20x and 40x magnification. To generate 20x magnification, we crop the same original domains inward by 27.5% in each direction to produce 368 × 368 µm 2-D slice images. 40x magnification is generated by cropping 38% in each direction to produce 184 × 184 µm 2-D slice images.

To study how z-gap size affects average void area fraction using our simulated particle domains, we sample z-slices within the middle 50% of the scaffold that are z-gap apart and plot average void area fraction as a function of z-gap. To study how the number of z-slices affects average void area fraction, we sample an increasing number of z-slices starting from the bottom of the scaffold that are 8 µm apart. For each iteration, we compute the average void area fraction and plot as a function of the number of z-slices. We use the same particle domains as described in the previous paragraph.

### Real granular scaffolds

We study how VVF measurements vary for four conditions: 1) scaffolds comprising 70 µm polyethylene glycol (PEG) vs. 100 µm hyaluronic acid (HA) particle scaffolds, 2) 10x, 20x, and 40x optical lens magnifications, 3) Fiji, MATLAB, Imaris, and 3-D Particle Segmentation software approaches, and 4) a reasonable range of input parameters or settings that produce a low, middle, and high VVF value. The goal of the last condition is to study the range of user bias for each software. As mentioned before, our protocol for determining reasonable low, middle, and high output images was as follows: low outputs showed many ‘false particle’ regions but at least one ‘false void’ region; middle output images showed both ‘false particle’ and ‘false void’ regions; and high output images showed many ‘false void’ regions but at least one ‘false particle’ region.

Hydrogel microparticles were generated in microfluidic devices, then interlinked to form MAP scaffolds. Samples were imaged using a Nikon C2 confocal microscope.

### Fiji

Fiji is an image processing package that approximates VVF by computing the average void area fraction among inputted 2-D z-slice images. For our analysis, we exclude blurry z-slices that appear out of focus. Additionally, we direct the program to use a slice-specific binarization threshold for each individual z-slice regardless of the thresholding method. To define our reasonable range of low, middle, and high outputs, we select three binary thresholding methods from Fiji’s 16 built-in Auto Threshold methods: MinError, Huang, and Default, respectively. These selections were based on the sample image of cells shown in the Fiji/ImageJ online documentation^10^, where we relate the circular cells to particles. MinError binarization correctly segments complete cells/particles but includes substantial particle-pixel noise, which underestimates void space. Huang produces the optimal cell/particle segmentation. Default binarization results in cells/particles that are speckled with void space pixels, which overestimates void space. These three methods reference actual settings used in the field for reporting VVF. In practice, the optimal threshold method depends on the image, but this experiment exemplifies how the method impacts the VVF measurement.

### MATLAB

Our MATLAB script uses standard built-in functions to process and binarize images, then approximates VVF by computing the average void area fraction among inputted 2-D z-slice images. Users input a parameter, γ, that divides the output of the MATLAB function graythresh (Otsu algorithm) used for binarization. We chose γ = 1 or 1.5 for our low output category, γ = 2 for our middle output category, and γ = 2.5 or 3 for our high output category. These γ values were chosen on a per-z-stack basis by observing how different γ values change the binarizations, then visually determining a ‘reasonable’ output for each category.

### Imaris

Imaris is a microscopy image analysis software that computes VVF by binarizing z-stacks before generating a triangulated mesh of solid regions and computing VVF from the resulting 3-D surfaces. Imaris requires a user-inputted ‘manual absolute intensity threshold’ parameter that is globally applied to all images in the z-stack, so differential light intensity among z-stacks will contribute to less accurate binarization. Light intensity cannot be easily normalized within z-slices or across z-slices in Imaris. Selecting threshold parameters for our low, middle, and high output categories was done on a per-z-stack basis.

### 3-D Particle Segmentation

The complete process for our 3-D Particle Segmentation approach includes intensity compensation, binarization, distance transform, and watershed. We first convert z-stacks into 3-D intensity images then pre-process the 3-D volume to reduce the data size and improve the image quality. We resize each 3-D volume according to a specified cubic voxel size, and we compensate for intensity variation over depth by subtracting each x-y plane with its 1% percentile. We binarize the 3-D volume by selecting foreground (tentative-particle) voxels using three sequential conditions: (1) The voxel intensity is larger than a minimum intensity, *I*_*min*_; (2) The voxel intensity is not lower than the mean of neighboring voxels by more than *I*_*diff*_. When particles are densely packed, the gap between particles is sometimes narrow, resulting in a void region that is brighter than *I*_*min*_. These void voxels are identified using relative intensity because they are significantly dimmer than neighboring voxels; (3) The voxel intensity is higher than a maximum intensity, *I*_*max*_. Because the relative brightness used in Condition (2) is based on absolute intensity difference, voxels with very high intensity may show high relative intensity variation that satisfies Condition (2). Therefore, we consider voxels with very high intensity as particle voxels, regardless if they satisfy Condition (2). Foreground (tentative-particle) voxels are marked as 1 if they satisfy Condition (3) or if they satisfy Conditions (1) and (2). We consider all other voxels as background (void space) voxels, marked as 0. To generate our low, middle, and high outputs, we adjust *I*_*min*_ on a per-z-stack basis, similar to our approach for MATLAB and Imaris. For the remaining parameters, we set *I*_*diff*_ = *I*_*min*_ / 3, and *I*_*max*_ = *I*_*min*_ * 333 based on reasonable test outcomes.

Next, we use watershed to segment the foreground voxels into discrete particles. We compute the 3-D distance transform of the volume, *D*, as the distance of each voxel to the nearest background voxel. We find the local maxima of *D* as the initial seeds. To avoid finding multiple seeds within the same particle, we identify close local maxima (defined as local maxima whose distance is smaller than the mean radius of the particles). If the intensities of the voxels between the close local maxima do not drop significantly relative to the local maxima, then we remove the lower local maximum and only keep the higher one. We then use 3-D watershed seeded with the remaining local maxima to segment the volume of *−D*.

Next, we refine the segmentation results by removing small discrete volumes. For each putative segmented particle from the previous step, we calculate the surface-to-volume ratio. If the ratio is larger than a threshold, we remove these ‘false’ particles along with their seeds. We then apply watershed to *−D* again using the remaining seeds, which allows some original ‘false’ particles to get absorbed into nearby ‘true’ particles. We recursively apply watershed to each resulting segmented volume, as long as the surface-to-volume ratios of the smaller volumes are smaller than the threshold. Finally, we smooth the segmented regions by removing sharp boundary voxels.

## Author contributions

LR designed the study, generated data for simulated scaffolds and plots for real scaffolds, analyzed results, created figures, and wrote the manuscript. GW developed the protocol used on real scaffolds, generated data for real scaffolds, generated real scaffold images for figures, and contributed to writing and editing the manuscript. YB developed the code for 3-D particle segmentation of real scaffolds, wrote the methods for his approach, and provided valuable feedback during manuscript revisions. PC generated all simulated particle domains and wrote the corresponding methods. KW and YL provided real scaffold images used in analysis and provided insight for analyzing results, and YL provided manuscript edits. YG provided input and direction for developing the 3-D particle segmentation code and provided valuable feedback during manuscript revisions. TS provided input and direction for designing the study, analyzing results, organizing figures, and guiding the project. All authors were given an opportunity to edit the manuscript.

## Acknowledgements

We would like to thank the National Institutes of Health, the National Institutes of Neurological Disorders and the Stroke (1R01NS112940, 1R01NS079691, R01NS094599), and the National Institute of Allergy and Infectious Disease (1R01AI152568).

